# Isolectin B4 (IB4)-conjugated streptavidin for the selective knockdown of proteins in IB4-positive (+) nociceptors

**DOI:** 10.1101/2023.12.18.572242

**Authors:** O Bogen, D Araldi, A Sucher, K Kober, PT Ohara, JD Levine

## Abstract

*In vivo* analysis of protein function in nociceptor subpopulations using antisense oligonucleotides and short interfering RNAs is limited by their non-selective cellular uptake. To address the need for selective transfection methods, we covalently linked isolectin B4 (IB4) to streptavidin and analyzed whether it could be used to study protein function in IB4(+)-nociceptors. Rats treated intrathecally with IB4-conjugated streptavidin complexed with biotinylated antisense oligonucleotides for protein kinase C epsilon (PKCε) mRNA were found to have: a) less PKCε in dorsal root ganglia (DRG), b) reduced PKCε expression in IB4(+) but not IB4(–) DRG neurons, and c) fewer transcripts of the PKCε gene in the DRG. This knockdown in PKCε expression in IB4(+) DRG neurons is sufficient to reverse hyperalgesic priming, a rodent model of chronic pain that is dependent on PKCε in IB4(+)-nociceptors. These results establish that IB4-streptavidin can be used to study protein function in a defined subpopulation of nociceptive C-fiber afferents.

## Introduction

Nociceptors are a heterogeneous group of sensory neurons specialized for the detection of potentially tissue damaging stimuli (1). Most nociceptors are either thinly myelinated Aδ- or unmyelinated C-fiber neurons (2). Based on differences in neurotrophin dependence and the expression of neuropeptides, C-fiber nociceptors have been further divided into nerve growth factor (NGF)-dependent, peptidergic and glial-derived neurotrophic factor (GDNF)-dependent, non-peptidergic, nociceptors (3). The latter class can, at least in rodents (4), also be characterized by binding to isolectin B4 (IB4) (5) (6), a homotetrameric carbohydrate-binding protein from the African shrub *Griffonia simplicifolia* with high binding specificity for terminal α-D galactopyranose residues (7) (8).

IB4(+)-nociceptors that have been exposed to an inflammatory or neuropathic insult develop long-lasting latent hypersensitivity to inflammatory mediators (9) (10) (11) (12). This phenomenon, hyperalgesic priming, can be demonstrated in behavioral experiments as an enhanced and prolonged hyperalgesic response to the pro-inflammatory cytokine prostaglandin E2 (PGE_2_) (13) (14). The associated neuroplasticity in the phenotype of IB4(+) nociceptors depends on protein kinase C epsilon (PKCε) and can be explained by a switch in the intracellular signaling pathway mediating PGE_2_ hyperalgesia (15) (16) (17) (18).

Binding of IB4 to α-D galactoside containing glycoconjugates on the plasma membrane of GDNF-dependent, non-peptidergic C-fiber nociceptors induces receptor-mediated endocytosis (19) (20). Various laboratories have used this finding to selectively ablate non-peptidergic C-fibers with IB4-conjugated saporin in order to investigate their functional contribution to pain (21) (22) (23) (24). Based on these studies, we sought to analyze whether this cellular uptake mechanism could be used for the specific transfection of GDNF-dependent, non-peptidergic nociceptors with antisense oligonucleotides. Here we used IB4-conjugated streptavidin to transfect non-peptidergic C-fibers with a biotinylated antisense oligonucleotide to PKCε mRNA and analyzed its effect on hyperalgesic priming, a well-characterized model for the transition from acute to chronic pain in rodents (13) (14).

## Material and Methods

### Animals

All experiments were performed on 260–320 g adult male Sprague–Dawley rats (Charles River Laboratories, Hollister, CA, USA). Animals were housed in a controlled environment (21-23°C, 12-hour day-night cycle, food and water were available *ad libitum*) in the animal care facility at the University of California, San Francisco. Experimental protocols were approved by the Institutional Animal Care and Use Committee (IACUC) at the University of California at San Francisco and in compliance with the National Institutes of Health *Guide for the care and use of laboratory animals*. Every effort was made to minimize the number of animals used in experiments and their suffering.

### Randall-Selitto paw-pressure test

Mechanical nociceptive threshold was quantified using an Ugo Basile Analgesymeter^®^ (Stoelting, Chicago, IL, USA), a device that uses a cone-shaped pusher with a rounded tip, to apply a linearly increasing mechanical force to the dorsum of a rat’s hind paw, as previously described (25) (26). Rats were lightly restrained in a cylindrical acrylic tube that has lateral ports at one end, to allow easy access to the hind paw for mechanical nociceptive threshold testing. Rats were acclimatized to the testing procedure by placing them in the restrainers and training them with the Analgesymeter for ∼2 h per day for 3 consecutive days before measuring baseline mechanical nociceptive threshold. Nociceptive threshold was defined as the force in grams at which a rat withdrew its paw. Baseline mechanical nociceptive threshold was defined as the mean of the three readings taken before test agents were injected.

### Drugs and their administration

The selective PKCε agonist, psi ε receptor for activated C kinase (ψεRACK) (27) was purchased from Biomatik (Wilmington, DE, USA), and the proinflammatory cytokine prostaglandin-E_2_ (PGE_2_) from Sigma-Aldrich (St. Louis, MO, USA). The selection of drug doses was based on dose-response relationships determined during earlier studies (28) (15). Stock solutions of PGE_2_ (1 µg/µl) in 100% ethanol (EtOH) were diluted (1:50) with physiological saline (0.9% NaCl) immediately before intradermal injection. The EtOH concentration of the final PGE_2_ solution was 2% (w/v) and the injection volume 5 µl. λφϑεRACK was dissolved and diluted in physiological saline (c_final_= 0.4 µg/µl). λφϑεRACK (1 µg) was administered by hypotonic shock as previously described (12). All drugs were administered via a 30-gauge beveled hypodermic needle connected to a Hamilton micro syringe with polyethylene tubing (PE-10).

### Covalent conjugation of streptavidin to IB4

The streptavidin conjugation kit (cat. #: ab 102921) was purchased from Abcam (Waltham, MA, USA) and IB4 (cat. #: L-1104-1) was purchased from Vector Laboratories (Burlingame, CA, USA). All reagents were equilibrated to room temperature (RT) before use. 100 μl of IB4 (c = 2.15 mg/ml), dissolved in sterile phosphate buffered saline (PBS), was activated by the addition of 20 μl of modifier reagent of the streptavidin conjugation kit, and then pipetted onto 100 μg of streptavidin (stochiometric ratio between IB4 and streptavidin = 1:1). This solution was gently mixed and incubated at room temperature for 24 h in the dark. Free excess amine groups on IB4 and streptavidin were inactivated by adding 20 μl of quenching reagent of the streptavidin conjugation kit to the reaction mixture. IB4-streptavidin conjugate was stored at 4°C until use.

### Intrathecal injection of antisense oligonucleotides

The antisense oligodeoxynucleotide (ODN) for PKCε, Biotin-GCO AGC TOG ATO TTG OGC OC (O = C-phosphorthioate) directed against a unique region of the rat mRNA was synthesized by Invitrogen (Waltham, MA, USA). The corresponding GenBank accession number and ODN position within the mRNA sequence are NM_017171.2 and 420-439, respectively. We have previously proven that this antisense ODN for PKCε can be used to downregulate PKCε expression in rat DRG (9) (26). The corresponding sense ODN sequence for PKCε, Biotin-GGE CGC AFG ATO GAG OTG EC (E = G-phosphorthioate, F = A-phosphorthioate), was used as the control. ODNs were dissolved in ultrapure water to a final concentration of 100 pmol/µl, aliquoted and stored at –20 °C, until use.

For each injection 2.3 μl IB4-streptavidin (30 pmol of conjugate with 120 pmol of biotin binding sites), 1.2 μl biotinylated ODN (c = 100 pmol/μl) and 16.5 μl PBS were mixed (injection volume = 20 μl), incubated at RT for 5 minutes and transferred into a 3/10 cc insulin syringe with a 29-gauge ultra-fine 1/2-inch fixed hypodermic needle (Walgreens, Deerfield, Il, USA). Rats were briefly anesthetized with 2.5% isoflurane (Phoenix Pharmaceuticals, St. Joseph, MO, USA) to facilitate intrathecal injection, and the hypodermic needle inserted into the subarachnoid space on the midline, between the L4 and L5 vertebrae (29). The intrathecal site of injection was confirmed by the elicitation of a tail flick produced by the insertion of the needle into the intrathecal space and injection of the solution (30). Intrathecal injections were performed on day 1, 4 and 7 and DRG harvested on day 10, to be used for either Western blotting, quantitative real-time PCR (Polymerase Chain Reaction) or immunohistochemical analysis.

### Tissue harvesting

Rats treated with antisense or sense ODN were euthanized by exsanguination while under deep isoflurane anesthesia. Then L4 and L5 DRG were surgically removed, snap frozen on dry ice and stored at –80° C until further processing.

### Protein extraction

DRG were transferred into homogenization buffer [150 mM NaCl, 50 mM Tris-HCl pH 7.4, 2 mM EDTA, 2% sodium dodecyl sulfate (SDS)] supplemented with complete protease inhibitor cocktail (Roche Diagnostics Corp., Indianapolis, IN, USA) and manually homogenized with a plastic pestle in an Eppendorf tube. Proteins were extracted by incubating the homogenate for 2 hours at RT in an Eppendorf thermomixer at 1400 rpm and separated from insoluble components by centrifugation for 15 minutes at 14,000 rpm in a tabletop centrifuge (Eppendorf, Hamburg, Germany). Protein concentration of the samples was determined using the micro-BCA protein assay kit (Pierce, Rockford, Il, USA) with BSA as a standard.

### Western blotting

Changes in PKCε expression were evaluated by Western blotting. Briefly, 20 μg of protein per sample were mixed with 4 x sample buffer (62.5 mM Tris-HCl pH 6.8, 3% SDS, 10% glycerol, 0.025% bromophenol blue), denatured by shaking for 10 min at 500 rpm and 90 °C in an Eppendorf thermomixer, and electrophoresed on a 4-15% precast polyacrylamide gel (Bio-Rad, Hercules, CA, USA) in 25 mM Tris buffer containing 192 mM glycine and 0.1% SDS. Proteins were transferred onto a nitrocellulose membrane (NC) using the semidry blotting method (transfer time 1 h at 10 V with 47.9 mM Tris, 38.9 mM glycine, 0.038% SDS and 20% methanol). The blotting membrane was saturated by shaking in Tris buffered saline (TBS), pH 7.4, containing 5% BSA and 0.1% Tween 20 (antibody dilution buffer) for 1 h at RT. The NC membrane was cut in half at ∼64 kDa and the upper half of the membrane was probed at 4°C overnight with a 1:500 dilution of a rabbit anti-PKCε antibody (sc-214, Santa Cruz Biotechnology, Dallas, Texas, USA) in antibody dilution buffer, while the lower half was probed at 4 °C overnight with a 1:1000 dilution of a rabbit anti-β actin antibody (ab8227, Abcam, Waltham, MA, USA), in antibody dilution buffer. Blots were rinsed with TBS containing 0.1% Tween 20 (3 times for 15 min under shaking at RT) and probed with an anti-rabbit HRP conjugated antibody (NA934V, GE healthcare, Piscataway, NJ, USA) for 2 h at RT. Blots were rinsed with TBS containing 0.1% Tween 20 (3 times for 15 min under shaking at RT) and PKCε and β-actin immunoreactivities were visualized with the enhanced chemiluminescence detection kit (Pierce) and analyzed by computer-assisted densitometry (31).

### Immunohistochemistry

Rats received 3 intrathecal injections of either IB4-streptavidin complexed with biotinylated antisense or sense oligonucleotides against PKCε mRNA on day 1, 4 and 7. On day 10 rats were deeply anesthetized with isoflurane and perfused through the left ventricle with 100 ml cold PBS containing 10 U heparin/ml PBS followed by 300 ml methanol-buffered 10% formalin (Thermo Fisher Scientific, Waltham, MA, USA). L4 and L5 DRG were dissected out and transferred to PBS with 30% sucrose and stored overnight at 4°C. The DRGs were then embedded in Tissue Tek (OCT compound, Electron Microscopy Sciences, Hatfield, PA, USA), sectioned in a Leica cryostat at 10 µm and mounted on gelatinized microscope slides (Southern Biotech, Birmingham, AL, USA).

Tissue Tek was removed by rinsing the DRG tissue sections twice with Tris-buffered saline (TBS) containing 0.1% Triton X-100 (TBST). Tissue sections were incubated for 1 hour at room temperature (RT) in TBST, with 10% normal goat serum (Jackson Immunoresearch, West Grove, PA, USA) (antibody dilution buffer). Sections were incubated overnight at 4°C with a 1:500 dilution of a Rabbit (Rb) anti-PKCε antibody (sc-214, Santa Cruz Biotechnology) in antibody dilution buffer, washed three times for 10 minutes each at RT with TBST and probed for 2h at RT with a 1:500 dilution of an Alexa Fluor conjugated Goat anti-Rb antibody (Invitrogen cat. #: A11012) and a 1:1000 dilution of IB4-FITC (Invitrogen, cat. #: I21411) in antibody dilution buffer supplemented with 0.1mM CaCl_2_, 0.1mM MnCl_2_, 0.1mM MgCl_2_ for 2h at RT (6). Sections were washed three times for 10 minutes at RT with TBST supplemented with 0.1mM CaCl_2_, 0.1mM MnCl_2_, 0.1mM MgCl_2_ and mounted with DAPI containing mounting media (Invitrogen, cat. #: P36935).

### RNA extraction and quantitative real-time PCR

Total RNA from L4 and L5 DRG was extracted using Trizol (Invitrogen) and the “PureLink^TM^ RNA Mini Kit” (Invitrogen) according to the manufacturer’s instruction. The RNA concentration in each sample was determined using a spectrophotometer (Shimadzu, Santa Clara, CA, USA). RNA was transcribed into cDNA using the “iScript^TM^ Advanced cDNA Synthesis Kit for RT-qPCR” (Bio-Rad, Hercules, CA, USA). The real-time PCR was performed with the “SsoAdvanced^TM^ Universal SYBR Green Supermix” (Bio-Rad) and specific primers for PKCε (Bio-Rad, assay ID: qRnoCID0008431) on the CFX 96^TM^ real-time PCR detection system (Bio-Rad). The PCR program was run under the following conditions: 2 min. at 95 °C, followed by 40 cycles of 15 s at 95 °C, and 30 s at 60 °C. Each sample was run in triplicate. Change in gene expression was determined by the 2^-1Δ1ΔCT^ method and expressed as relative change compared to control level (32). Glycerinaldehyde-3-phosphate dehydrogenase (GAPDH) RNA was used as reference gene (Bio-Rad, assay ID: qRnoCID0057018).

### Statistical analysis

a) Western blot and quantitative real-time PCR: Statistical comparisons were made with GraphPad Prism 10 statistical software (GraphPad Software In., La Jolla, CA, USA). Comparison between treatments were performed using Student’s t-test and a p-value less than 0.05 was considered statistically significant. b) Immunohistochemistry: Statistical comparisons were made with GraphPad Prism 10 statistical software. Comparisons between treatments were performed using contingency testing and Fisher’s exact test. A p-value of less than 0.05 was considered statistically significant. c) Behavior: Statistical comparisons were made with GraphPad Prism 10 statistical software. Comparisons between treatments were performed using two-way repeated measures ANOVA with *Post hoc* analysis. A p-value of less than 0.05 was considered statistically significant.

### Data availability

The data generated in this study are available upon request from the corresponding author.

## Results

### Transfection with IB4-streptavidin

To evaluate whether IB4-streptavidin can be used as a transfection reagent, rats were treated with a complex of IB4-streptavidin and biotinylated antisense ODN (or for the control biotinylated sense ODN) directed against PKCε mRNA. Two groups of 3 rats each received three intrathecal injections on days 1, 4 and 7 before their L4 and L5 DRG were harvested on day 10. Western blot analysis of protein extracts from DRG from the group that had been subjected to the antisense ODN treatment showed a significant decrease (∼40%) in the PKCε expression (9.5 +/-1.2 a.u., [N=3]) compared to the group that was treated with the sense ODN sequence (15.9 +/- 1.3 a.u., [N=3], Student’s t-test, P = 0.01, **Fig. 1A, B**) indicating successful knockdown of PKCε.

**Figure 1:**
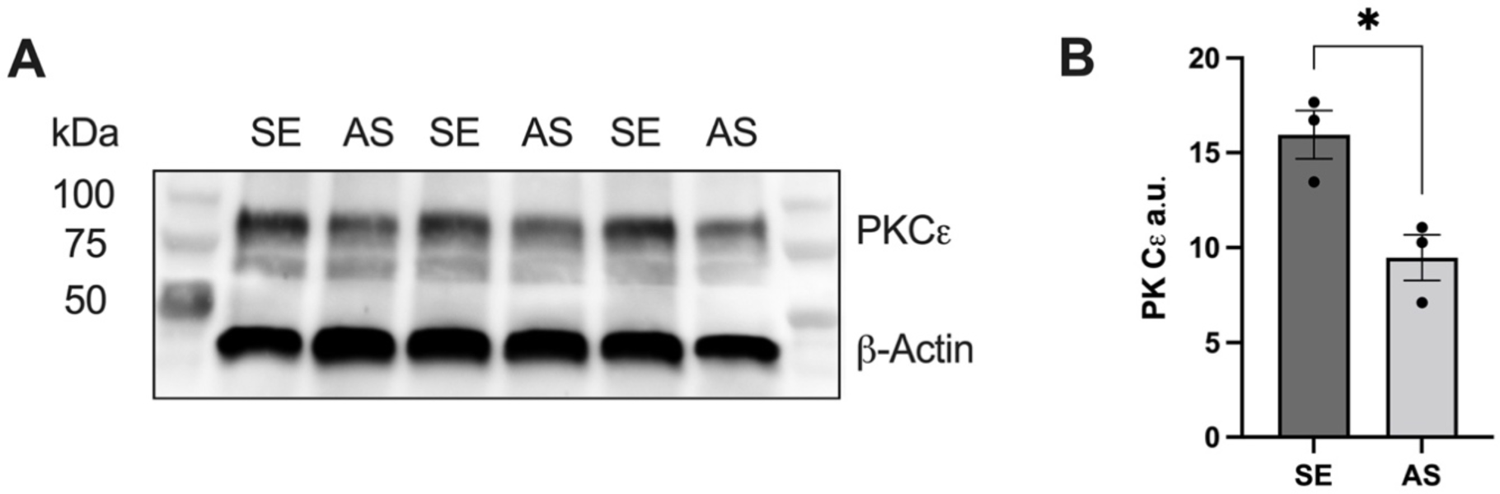
Effect of intrathecal PKC: antisense oligonucleotides on PKC: expression in rat lumbar dorsal root ganglia. (A) Western blot image of protein extracts from L4/L5 dorsal root ganglia of rats treated with either biotinylated PKC: sense (SE) or antisense (AS) oligonucleotides (bound to IB4 streptavidin). The calculated molecular weight of PKC: is ∼83.5 kDa (according to UniProt database entry P09216). β-Actin was used as a loading control. Its calculated molecular weight is ∼42 kDa (according to UniProt database entry P60711). (B) Column bar graph showing normalized PKC: immunoreactivity in both groups. Data are presented as mean +/- SEM with N=3 in each group (P<0.05).

### Immunohistochemistry

To analyze whether the downregulation of PKCε expression is restricted to IB4-positive DRG neurons, an immunohistochemical analysis was performed. Two groups of 3 rats each received three intrathecal injections of either biotinylated PKCε sense oligonucleotides bound to IB4-streptavidin or biotinylated PKCε antisense oligonucleotides bound to IB4-streptavidin on days 1, 4, and 7. Three days later, rats were transcardially perfused with 4% PFA, their L4 and L5 DRG excised and analyzed for PKCε immunoreactivity (**Fig. 2**). A contingency test was performed to compare the proportion of IB4(+)/PKCε(+) neurons between DRGs from rats treated with the IB4-streptavidin biotinylated-PKCε sense ODN complex and rats treated with the IB4-streptavidin biotinylated PKCε antisense ODN complex (IB4(+)/PKCε (+) in PKCε-sense ODN-treated rats [31%, 191/600 neurons, 95% CI = (28-35)%], in PKCε antisense ODN-treated rats [19%, 117/600 neurons, 95% CI = (16-22)%]). Fisher’s exact test shows a significant difference between IB4(+)/PKCε(+) neurons in DRG of rats treated with IB4-streptavidin biotinylated PKCε sense ODN and those treated with IB4-streptavidin biotinylated PKCε antisense ODN [P < 0.0001, N = 600/group (3 rats per group with 200 neurons/DRG per rat; 95% CI were calculated using the Wilson/Brown method)]. A second contingency test was performed to compare the proportion of IB4(-)/PKCε(+) neurons between DRGs from rats treated with the biotinylated PKCε sense ODN IB4-streptavidin complex and rats that were treated with the biotinylated PKCε antisense ODN IB4-streptavidin complex (IB4(-)/PKCε (+) in PKCε sense ODN-treated rats [10%, 65/600 neurons, 95% CI = (8-13)%], in rats treated with PKCε antisense ODN [11%, 70/600 neurons, 95% CI = (9-14)%]). Fisher’s exact test shows no significant difference between IB4(-)/PKCε(+) neurons in DRG of rats treated with IB4-streptavidin bound to biotinylated PKCε sense ODN and those treated with IB4-streptavidin bound to biotinylated PKCε antisense ODN [P=0.7, N=600/group (3 rats per group with 200 neurons/DRG per rat; 95% CI were calculated using the Wilson/Brown method). These results demonstrate that the knockdown in PKCε expression is limited to IB4-positive (+) neurons, most of which are known to be non-peptidergic C-fiber nociceptors (33) (34).

**Figure 2:**
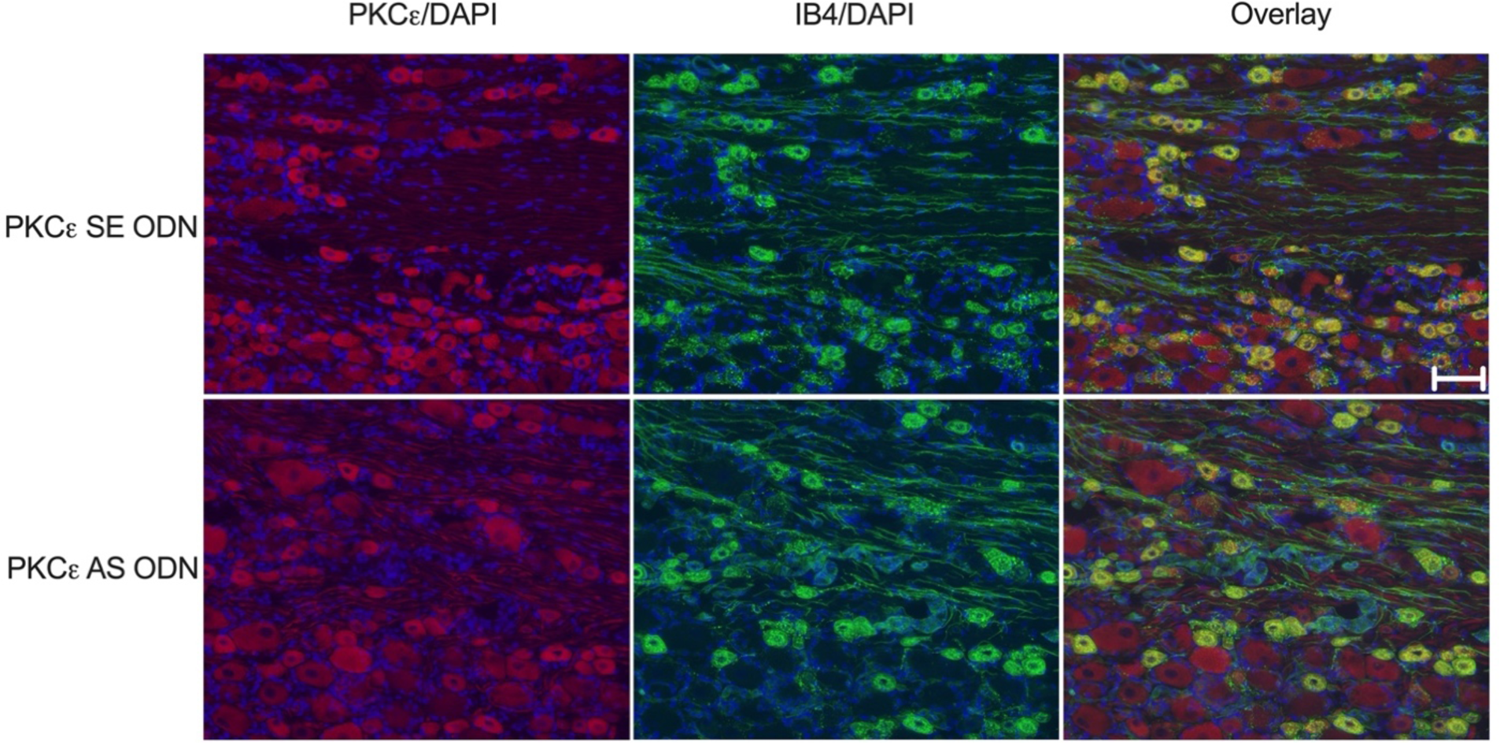
Knockdown of PKCε in IB4(+) C-fiber afferents. Representative images from L4/L5 DRG from rats treated with either IB4-streptavidin biotinylated PKCε sense ODN (upper image panels) or IB4-streptavidin biotinylated PKCε antisense ODN (lower image panels). Color coding: Red = PKCε immunoreactivity, Blue = DAPI, Green = IB4 immunoreactivity, Yellow = PKCε/IB4 colocalization. Note that up to 40% of all DRG neurons in the rat are IB4 positive (+) and the vast majority are small diameter C-fiber DRG neurons. PKCε immunoreactivity can be observed in small-, medium-, and large-diameter DRG neurons. Interestingly, the vast majority of PKCε expressing DRG (∼90%) are IB4 positive (+). Scale bar = 50 µm.

### Quantitative real-time PCR

To determine whether the decrease in PKCε protein levels in rat DRG is due to antisense ODN-stimulated degradation of transcripts of the PKCε gene or to a reduction in the frequency with which the PKCε mRNAs are translated into the corresponding amino acid sequence we performed a real-time PCR. Rats received three intrathecal injections with either IB4-streptavidin complexed with biotinylated antisense oligonucleotides to PKCε, or IB4-streptavidin complexed with biotinylated sense oligonucleotides to PKCε, on days 1, 4 and 7 before their L4 and L5 DRG were harvested on day 10. Quantitative real-time PCR analysis of RNA extracts from DRG from rats subjected to the antisense treatment showed a significant decrease in the number of PKCε transcripts (24% +/- 2.3%, [N=6], P<0.0001, **Fig. 3**) compared to the group of rats treated with the sense sequence, suggesting that the reduction in cellular PKCε protein levels is caused by a reduction in the number of transcripts of the PKCε gene in rat DRGs.

**Figure 3:**
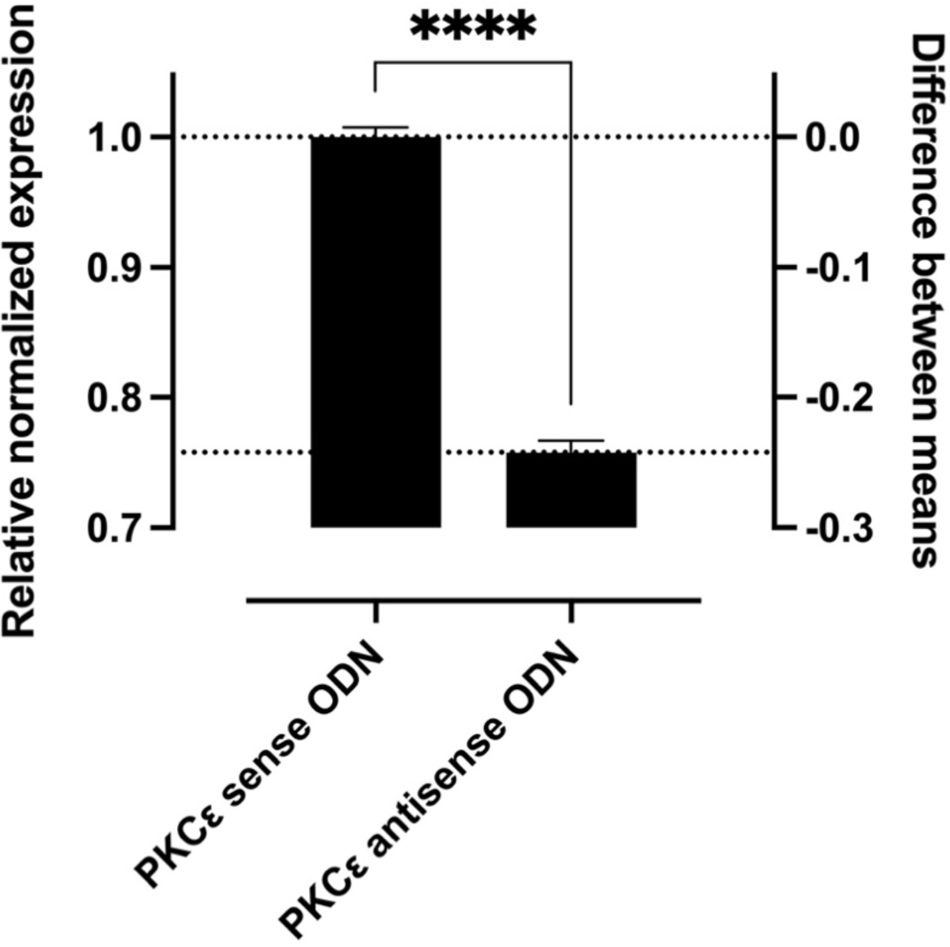
Effect of intrathecal PKCε antisense oligonucleotides on PKCε transcripts in rat L4 and L5 DRG. The histogram depicts the level of PKCε expression in rats treated with either a complex of IB4streptavidin/biotinylated PKCε sense ODN (left) or IB4 streptavidin/biotinylated antisense ODN (right). Gene expression was determined with SYBR green RT-PCR and values (mean +/- SEM) represent expression relative to GAPDH. Rats treated with IB4 streptavidin/biotinylated PKCε antisense ODN show a significant decrease in their PKCε mRNA levels compared to those treated with a mixture of IB4 streptavidin/biotinylated PKCε sense ODN (Student’s t-test, P=0.0001, N=6 for each group).

### Reversal of hyperalgesic priming in IB4(+)-nociceptors

Finally, to assess the potential therapeutic utility of our transfection method, we tested the efficacy of the knockdown on hyperalgesic priming, a well-characterized preclinical model of chronic pain (13) known to be dependent on PKCε activity in IB4(+)-nociceptors (11) (10). Rats were first primed with the selective PKCε activator \+JERACK (15), then treated with the complex of IB4-streptavidin and biotinylated PKCε antisense ODN (or biotinylated PKCε sense ODN for the control), and finally tested for the presence of hyperalgesic priming. Rats treated with the IB4-streptavidin biotinylated PKCε sense ODN complex demonstrate prolonged PGE_2_ hyperalgesia, a behavioral phenotype that characterizes the presence of hyperalgesic priming (**Fig. 4**) (9) (13). Rats treated with the IB4-streptavidin biotinylated PKCε antisense ODN complex only show the short lasting PGE_2_ hyperalgesia, a behavioral phenotype that characterizes the wild-type nociceptor (9) (13). These results demonstrate that the knockdown of PKCε in IB4(+)-DRG neurons can reverse the neuroplastic changes associated with the primed IB4(+)-nociceptor.

**Figure 4:**
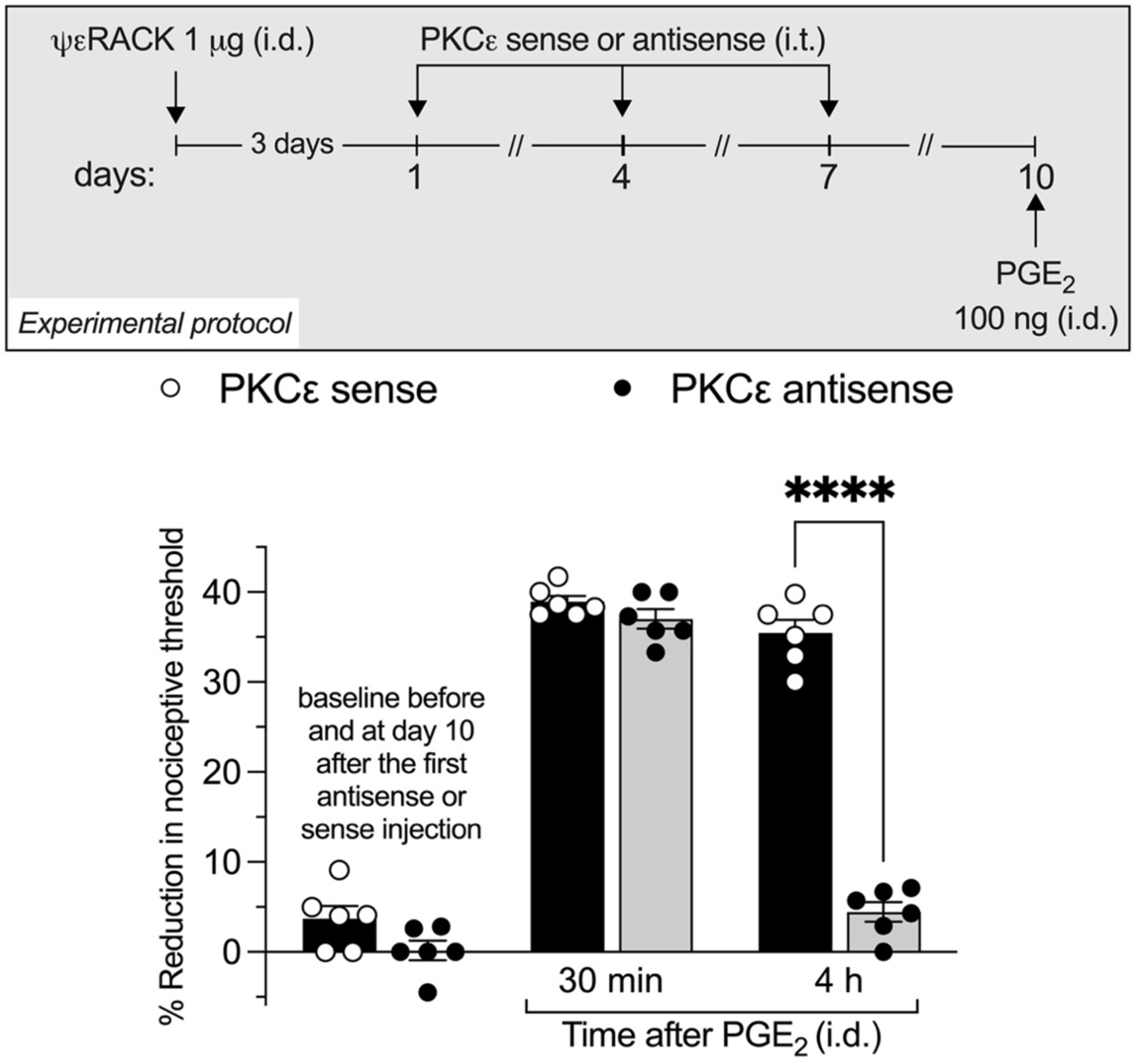
Reversal of hyperalgesic priming: Rats were primed by the intradermal administration of ψε:RACK. Starting three days after the injection of ψε:RACK, rats received intrathecal injections on day 1, 4, and 7 of IB4-streptavidin complexed with either biotinylated sense or antisense ODN for PKC: mRNA. Three days after the last injection PGE_2_ was administered into the same site asψε:RACK before. To test for hyperalgesic priming nociceptive paw withdrawal thresholds were measured prior to and 30 min and 4 h post intradermal administration of PGE_2_. PGE_2_-induced hyperalgesia remained elevated at 4h in sense-treated rats, indicating the presence of priming. However, hyperalgesia was not present at the 4^th^ h in antisense-treated rats, showing its ability to reverse priming. A two-way repeated measures ANOVA for the baseline readings showed no significant effect of treatment (F_1,10_ = 0, P > 0.999), time (F_2_,_20_ = 1.25, P = 0.308) and treatment by time interaction (F_2_,_20_ = 1.67, P = 0.225). These results show that the two ODN groups did not differ significantly until the PGE_2_ injection. A second two-way repeated measures ANOVA that included the two post-PGE_2_ injection time points showed significant effects for treatment (F_1_,_10_ = 159, P < 0.0001), time (F_1_,_10_ = 426.2, P < 0.0001) and treatment by time interaction (F_1_,_10_ = 277.9, P < 0.0001). *Post hoc* analysis revealed significant differences between sense and antisense treated rats only 4 h post PGE_2_ administration (t_20_= 19.74, P < 0.0001).

## Discussion

Opioids are the most effective analgesics for moderate to severe (acute, perioperative, posttraumatic and chronic) pain (35). However, their many undesirable side effects such as analgesic tolerance and physical dependence, opioid-induced hyperalgesia and respiratory depression mandate the development of novel analgesics (36). Therapeutic oligonucleotides are a possible solution (37) (38) (39) (40). Antisense oligonucleotides, or small interfering RNAs, are potent and highly specific (41) (42) (43), safe (41) (42) (43), their mode of action is understood (44) (45) (46) (47), and their *in vivo* stability has been significantly increased in recent years through chemical modifications (48) (45) (41) (42) (43). In addition, their use has already contributed to a better understanding of many pain syndromes (49) (50) (18) (51) (52) (53) (54). One of the problems still to be solved is how to limit the uptake of the oligonucleotides to specific cell types (55) (44) (46). The most promising method to date involves the conjugation of oligonucleotides to ligands known to bind to cell surface molecules that are exclusively expressed by the target cells and whose binding is known to trigger the cellular uptake of the ligand-conjugated oligonucleotide via receptor-mediated endocytosis (56) (57) (58) (59) (60) (61). Here we have used this approach to transfect a subpopulation of C-fiber nociceptors implicated in specific pain syndromes (e.g., oxaliplatin-induced painful peripheral neuropathy (62) and spinal nerve ligation (63)) with antisense oligonucleotides for PKCε mRNA. Our Western blot data show that a molecular conjugate of IB4-streptavidin complexed with biotinylated PKCε antisense oligonucleotides can be used to attenuate levels of PKCε in the rat dorsal root ganglia (**Fig. 1**). This result is consistent with the findings that IB4 binds to glycosylated cell surface molecules at a large number of small diameter sensory neurons (64) (65) (66), streptavidin is a biotin binding molecule (67) (68), and the antisense sequence we used is known to effectively downregulate PKCε expression in rat DRG (9) (26). Our immunohistological data show that the downregulation of PKCε is restricted to IB4-binding neurons (**Fig. 2**), most of which are known to be non-peptidergic C-fiber nociceptors (33) (34). This result is consistent with findings that IB4 binding to α-D-galactosides induces cellular uptake of the “ligand-receptor complex” through receptor-mediated endocytosis (19) (20). Our real-time PCR results show that the number of transcripts of the PKCε gene in RNA samples from DRG from rats treated with antisense oligonucleotides is lower than in RNA samples from DRG from rats treated with the sense oligonucleotides (**Fig. 3**). This suggests that the attenuation of PKCε expression in rats treated with antisense oligonucleotides is due to an RNase H-catalyzed degradation of PKCε mRNA (69) (70). Finally, our behavioral data show that the knockdown of the PKCε expression in IB4-binding neurons can reverse hyperalgesic priming (**Fig. 4**), a chronic pain phenotype known to be mediated by PKCε activity in non-peptidergic C-fiber nociceptors (9) (10) (12). This demonstrates that ligand-conjugated streptavidin can be used to selectively transfect target cells with biotinylated antisense oligonucleotides, and a similar approach using a lectin- or antibody-conjugated streptavidin could also be used for the regulation of proteins in other subpopulations of nociceptors.

Here, we used streptavidin instead of avidin to conjugate with IB4. The rationale for this is, that others have previously used streptavidin-conjugated ligands (56) and antibodies (71) to successfully transfect cells with biotinylated oligonucleotides, and the binding of biotin to streptavidin is pH dependent (67), while the binding of biotin to avidin is itself surprisingly stable under acidic and alkaline conditions (67) (68). One of the major challenges in developing oligonucleotide-based therapies is solving the problem of endosomal escape (48) (72) (73). All oligonucleotides, whether unconjugated, bound to plasma proteins, or as part of a ligand-conjugated complex, are taken up by cells through endocytosis and thus enter membrane-bound vesicles from which they must escape to fulfill their function. The mechanisms by which oligonucleotides escape from endosomes are still not fully understood (72) (74). However, it is known that the internal milieu of endosomes acidifies over time, which, for example, contributes to the breakdown of ligand-receptor complexes into their components. Given that the affinity of streptavidin for biotin decreases dramatically with decreasing pH (67), we chose to bind IB4 to streptavidin.

Ligand-conjugated oligonucleotides offer several significant advantages over unconjugated oligonucleotides. They have a much higher selectivity and are taken up by cells much more effectively (55) (44) (46). Both factors not only reduce the risk of unwanted side effects, but also allow a significant reduction in the doses needed for a successful knockdown. For example, in this study we used only intrathecal injections of 0.78 µg antisense ODN/dose, while our laboratory typically uses intrathecal injections of between 40 and 120 µg antisense ODN/dose for knockdowns with comparable potency (75) (76). However, a fundamental problem in using ligand-conjugated oligonucleotides is the identification of cell surface molecules that are specific to the target cells since most cell surface molecules are not expressed by only one cell type. This also explains why the development of ligand-conjugated oligonucleotides is so challenging and so far, only a few drugs based on ligand-conjugated oligonucleotides have been approved by the FDA (42) (77) (78).

In summary, we report the successful use of streptavidin-conjugated IB4 to selectively transfect a subset of small-diameter nociceptors, the IB4-binding C-fibers, with biotinylated antisense oligonucleotides for PKCε. Our results provide proof of concept for the regulation of target molecules in other populations of DRG neurons.

## Acknowledgements

This work was supported by funding from the National Institutes of Health (NIH) ([1] R01AR075334; [2] NIH R01CA250017). The NIH had no role in study design, data collection and interpretation, or the decision to submit the work for publication. Furthermore, the content is solely the responsibility of the authors and does not necessarily reflect the official views of the NIH.

## Notes

### Competing Interest Statement

The authors have declared no competing interest.

### Summary of Updates

1) Data availability 2) Source of funding (NIH)

